# Disentangling mucus rheology and transport efficiency in human airways

**DOI:** 10.64898/2026.01.10.698668

**Authors:** Alice Briole, Qian Mao, Umberto D’Ortona, Julien Favier, Annie Viallat, Etienne Loiseau

## Abstract

The protection of the respiratory tract relies on a layer of mucus that is transported along the epithelial surface by the beating of millions of microscopic cilia. This mechanism, called mucociliary clearance, is defective in chronic respiratory diseases. Since these pathologies alter mucus rheology, mucus transport efficiency is hypothesized to rely on its mechanical properties. Yet, this link is difficult to test due to limited experimental models. Here, we introduce an experimental framework that enables us to identify conditions associated with efficient mucus transport or arrest. Strikingly, we show that cilia are able to efficiently propel mucus with properties ranging from a viscoelastic fluid to an elastic solid, revealing that bulk mucus rheology is not the critical determinant of clearance failure. Instead, we demonstrate that the efficiency of mucociliary clearance is governed by the hydration of a thin fluid layer at the cilia-mucus interface. Finally, we use a hydrodynamic model informed by measurements on ciliary beat patterns to infer the properties of this critical layer under both transported and arrested mucus conditions. This work not only offers a novel understanding of the fundamental physical mechanism of mucus transport, but also provides a well-defined and quantitative assay to test effects of mucolytic agents or drugs on mucus clearance.

## 1 Introduction

Mucociliary clearance is the primary defense mechanism of the airways. It consists of a protective layer of mucus, lining the epithelial surface, transported by the continuous beating of millions of microscopic cilia. This system is essential for clearing inhaled pathogens and particulate matter from the lungs.

Chronic respiratory diseases such as severe asthma, chronic obstructive pulmonary disease (COPD), and cystic fibrosis are all associated with defective clearance [1–3]. In these pathologies, mucus production and composition change significantly, often involving hypersecretion, altered mucin expression, and the presence of immune cells, DNA, and bacteria following infection [4–6]. Studies performed on patient-derived sputum samples have shown that these compositional changes impact mucus rheology [7–9] and have aimed to establish links between rheological properties and disease severity [10, 11]. As a consequence, it is widely assumed that mucus transport efficiency is primarily governed by its mechanical properties, with increased viscosity and/or elasticity impairing mucociliary clearance, up to the extreme case of airway obstruction by mucus plugs.

However, airway mucus is a heterogeneous material, making quantitative rheological measurements difficult to interpret, particularly when sample collection and preparation are not carefully controlled [5]. To overcome some of the limitations intrinsic to sputum handling, *in vitro* models of human bronchial epithelium grown at the air–liquid interface (ALI) provide a promising alternative to study airway mucus. The small volumes produced at the surface of such cultures are well adapted to microrheology techniques, enabling the characterization of mucus microstructure, heterogeneities, and spatial variations of rheological properties along the height of the mucus layer [4, 12–15]. Yet, it remains unclear whether rheological measurements performed on ALI cultures can be directly interpreted as representative of pathological airway mucus observed in respiratory diseases [13, 16]. Moreover, studies that directly link rheological properties to mucus transport efficiency remain scarce [11, 17, 18]. While Ma et al. reported a correlation between increased elastic and viscous moduli of cystic fibrosis sputum and reduced transport [11], Roy et al. highlighted the necessity of MUC5B expression for efficient transport [17]. As a result, testing the widely shared hypothesis that mucus rheology directly governs transport efficiency remains challenging, as most experimental studies focus either on mucus rheology or on cilia dynamics [19–21].

Numerical modeling provides a powerful framework to get new insights on physical mechanisms governing mucociliary clearance. While a vast corpus has explored ciliary systems [22–24], most numerical studies rely on prescribed ciliary beat patterns [25–28]. These approaches imply a one-way coupling that neglects the essential fluid feedback on ciliary deformation, thereby limiting their ability to capture the system’s dynamic response to changing rheological environments. To achieve a more realistic description, three-dimensional models incorporating two-way coupling between the cilia and the two-phase airway surface flow (periciliary layer and mucus) have recently been developed [29]. However, they are rarely informed by quantitative measurements done on ciliary epithelia.

Here, we use a well-characterized Air-Liquid Interface (ALI) model that forms large-scale organized mucus vortices [30] to investigate this exact relationship. By combining transport measurements with *in situ* microrheology, we show that cilia are able to efficiently propel mucus with properties ranging from a viscoelastic fluid to an elastic solid. We challenge the paradigm that bulk mucus rheology governs transport and instead provide evidence for a critical layer at the cilia-mucus interface by proposing a new model where transport properties rely on the hydration state of this interface. Finally, we use computational fluid dynamics to model the cilia-mucus hydrodynamic interaction. This model, informed by measurements on ciliary beat patterns, enables us to infer properties of the interfacial layer for both transported and arrested mucus conditions.

## 2 Results

We used bronchial epithelial cultures at Air-Liquid Interface (ALI), derived from human primary cells of healthy donors (**Figure 1A**). In these cultures, the basal side is in contact with the culture medium through a porous membrane while the apical surface is exposed to air. These cultures differentiate to recapitulate the pseudostratified mucociliary epithelium phenotype [31, 32], featuring mucus-producing goblet cells and ciliated cells. Each ciliated cell carries 200–300 active cilia [33], which are grouped into bundles.

**Fig. 1.**
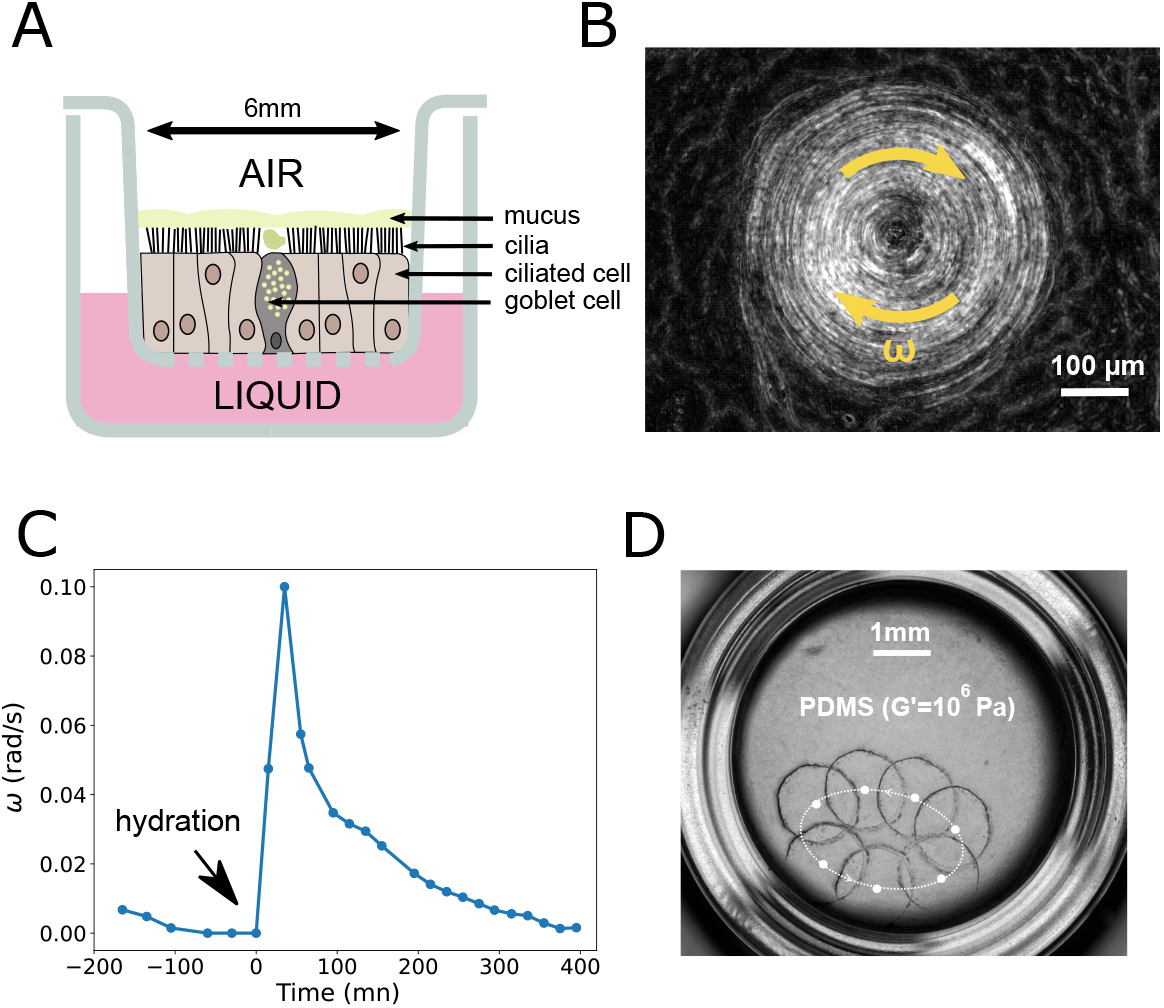
Observations pointing to an interfacial failure. (A) Schematic of the *in vitro* (ALI) model. (B) Representative image of a stable mucus vortex. The image is the result of a standard deviation projection over 300 frames acquired at 25 fps. (C) The angular velocity (*ω*) of a native vortex slows down and then restarts almost instantaneously upon apical addition of PBS (hydration). (D) Microscopy image showing the active transport of a piece of PDMS, an elastomer whose elastic modulus *G*′ ≈ 10^6^ Pa. The PDMS was added on a mucus-free surface.

In this system, hydrodynamic coupling between beating cilia and mucus results in the formation of deterministic fluid flow patterns that form macroscopic mucus vortices [30, 34, 35] (**Figure 1B**). In the following, we take advantage of these stable and well-defined structures to quantify the mucus transport and investigate the conditions associated with efficient transport or mucus arrest.

### Evidence for an interfacial failure mechanism

A first key observation on ALI culture is that the velocity of mucus transport is extremely sensitive to hydration. Indeed, the angular velocity of a mucus vortex, maintained in a humidity saturated environment, slows down over several hours till a complete arrest (**Figure 1C**). Strikingly, the addition on the apical surface of 1.5 µl of PBS results in a rapid recovery of transport within seconds. Although water diffuses rapidly (*D*_*w*_ ≈ 10^−9^ m^2^/s [36]), gel swelling is limited by the relaxation of the polymer network and is therefore governed by an effective collective diffusion coefficient *D*_*p*_ [37], typically 10^−11^–10^−12^ m^2^/s for biological gels [38, 39]. For a mucus layer of thickness *h* = 75 µm, this yields characteristic swelling times (*τ* = *h*^2^/*D*_*p*_) of 10^3^–10^4^ s (10 minutes to over an hour). The near-instantaneous recovery of transport upon PBS addition therefore cannot arise from bulk swelling, but instead indicates a direct interfacial effect.

To further support the hypothesis that the hydration state of the cilia-mucus interface is a critical parameter governing mucus transport properties — outweighing the potential role of the bulk rheology of mucus — we performed the following experiment. We selected a cell culture that exhibited stable mucus transport. We removed the native mucus by washing the apical surface with Dithiothreitol (DTT), followed by three successive rinses with PBS. We placed on the culture a small piece of cross-linked PDMS, which has a high elastic modulus (*G*′ ≈ 10^6^ Pa), many orders of magnitude stiffer than mucus (*G*′ ≈ 1 − 10 Pa [7, 16, 40]). While the piece of PDMS was initially immobile, a slight re-hydration (1.5 µL PBS) induced a rotational movement of the solid-elastic object (**Figure 1D** and **Supplementary Video S1**). This movement continued until the PDMS eventually adhered to the edge of the culture insert.

The observation that a solid-elastic object such as PDMS (*G*′ ≈ 10^6^ Pa) is transported at velocities comparable to those of native mucus (*G*′ ≈ 1 Pa) provides strong evidence that bulk rheological properties are not the primary determinant of transport. Instead, the cilia-mucus interface appears to play a crucial role. To get better insight on the nature of this interface and the fundamental mechanisms at play to efficiently transport mucus, we proceed in three steps. First we address the lack of a common standard for transport measurement in ALI culture by establishing a standardized metric. Second, we perform *in situ* microrheology measurements and correlate them with transport properties. This way, we rigorously test the role of mucus rheology. Finally, we quantify cilia beat pattern and use it as a physical probe to investigate the properties of the cilia-mucus interface.

### Standardizing mucus transport measurement

Quantification of mucus transport on ALI cultures lacks established methods to compare transport properties between different samples. To overcome this limitation, we propose to use cell cultures that exhibit robust mucus vortices. They provide a controlled experimental platform to assess mucus transport. First, it offers a well-defined, organized area accounting for the underlying ciliary organization. Second, we observed that mucus, within these vortices, moves as a solid body; debris (such as dead cells or added beads) shows no relative motion (**Supplementary Video S2**). This simplifies the transport field to a single angular velocity (*ω*), rather than a complex map of local velocities.

To compare vortices with different radius, R, and mucus thickness, h, we introduce the volumetric flow rate, Q. Based on the solid-body rotation (which implies *v*(*r*) = *ωr*), the flow rate is derived by integrating the velocity field over the radial crosssection, yielding 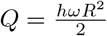 (**Figure 2A**).

**Fig. 2.**
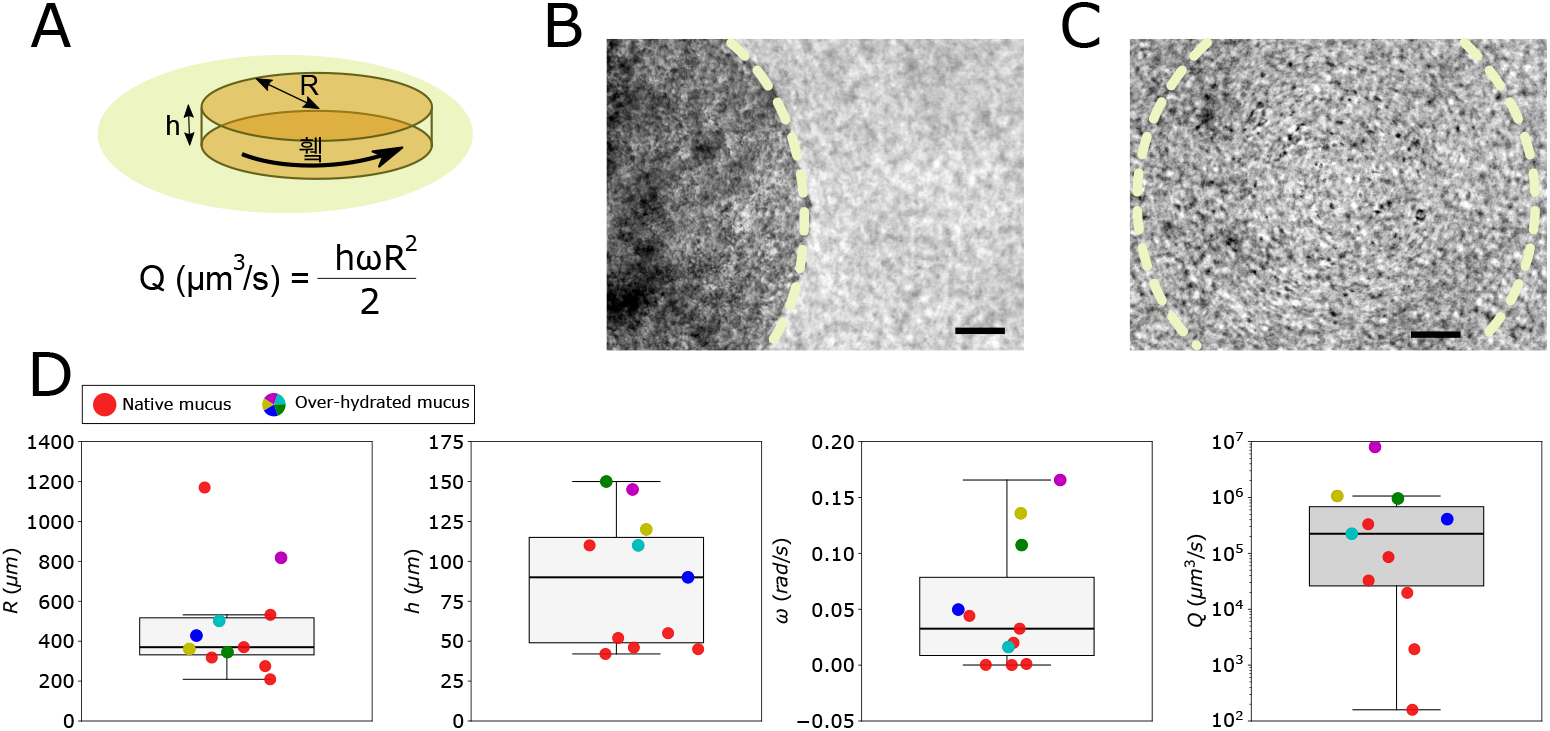
Standardizing *in vitro* transport quantification. (A) Schematic of a mucus vortex characterized by its radius (*R*), height (*h*), and angular velocity (*ω*). The volumetric flow rate 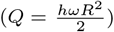 is used as a standardized metric for transport efficiency. (B) Bright-field image of a native, concentrated mucus vortex. (C) Bright-field image of an over-hydrated mucus vortex. The dotted lines indicate the vortex boundaries. Scale bars: 50 µm. (D) Box plots showing the distribution of measured vortex parameters: Radius (R), height (h), angular velocity (*ω*), and calculated flow rate (Q). In this study, N=11 vortices.

We applied this metric to N=11 distinct vortices, each obtained from a different cell culture. Our dataset presents two distinct experimental conditions: 6 “native” vortices and 5 “over-hydrated” ones. We over-hydrated the mucus by repeated addition of PBS on the apical surface to modify its initial rheological properties. The difference in mucus aspect between these two conditions is illustrated in **Figures 2B** and **2C** (see also **Supplementary Video S3**).

The vortex parameters are summarized in **Figure 2D**. The geometrical parameters show significant, yet expected, variability. The vortex radius *R* represents the characteristic length scale over which mucus is transported in a coordinated way. The size of the self-organized domain depends on cilia density and the frequency of apical washes. The mucus thickness *h* varies as the mucus production rates differ between cultures and is also affected by recent apical washes. The median value (*h* = 75 µm) is substantially greater than the ∼ 10 µm layer found in human bronchi [41]. This can be explained by mucus accumulation in ALI systems.

Calculated flow rates (Q) for both native and diluted mucus span several orders of magnitude (10^2^ − 10^7^ µm^3^/s). Importantly, the estimated physiological flow rate in human airways (calculated for bronchioles, *Q*_physio_ = (2*πRh*) × *v* using *R* = 500 µm, *h* = 7 − 10 µm, and *v* = 3 − 10 µm/s [42]) is ≈ 1 − 2 × 10^5^ µm^3^/s, which falls well within our experimentally observed range. In the following, the dilution of mucus enables us to correlate transport efficiency with rheology across a wide range.

### Linking mucus rheology and transport properties

Having established a standardized transport metric (Q), we detail our method for *in situ* microrheology measurement. We then quantitatively test the relationship between mucus rheology and transport efficiency.

### *In situ* microrheology of mucus

The typical volume of mucus per vortex is in the order of 40 nL (*V* = *πR*^2^*h* with R ∼ 400 µm and h ∼ 75 µm). Passive microrheology is a well-suited method to probe local rheological in small volumes. It relies on tracking Brownian particles [43–45]. The primary challenge in our system is that active ciliary beating masks the Brownian motion. We thus developed a method to stop ciliary activity based on findings that cinnamaldehyde reversibly inhibits ATP production - necessary for cilia beating - by targeting mitochondria [46]. The addition of 22.5 mM cinnamaldehyde to the basal medium induces a decrease of ciliary beat frequency (CBF) till a complete cessation of ciliary activity after ∼ 40 minutes (**Figure 3A** and **Supplementary Video S4**). This approach enabled us to perform in-situ passive microrheology on the mucus layer immediately after quantifying its transport properties.

**Fig. 3.**
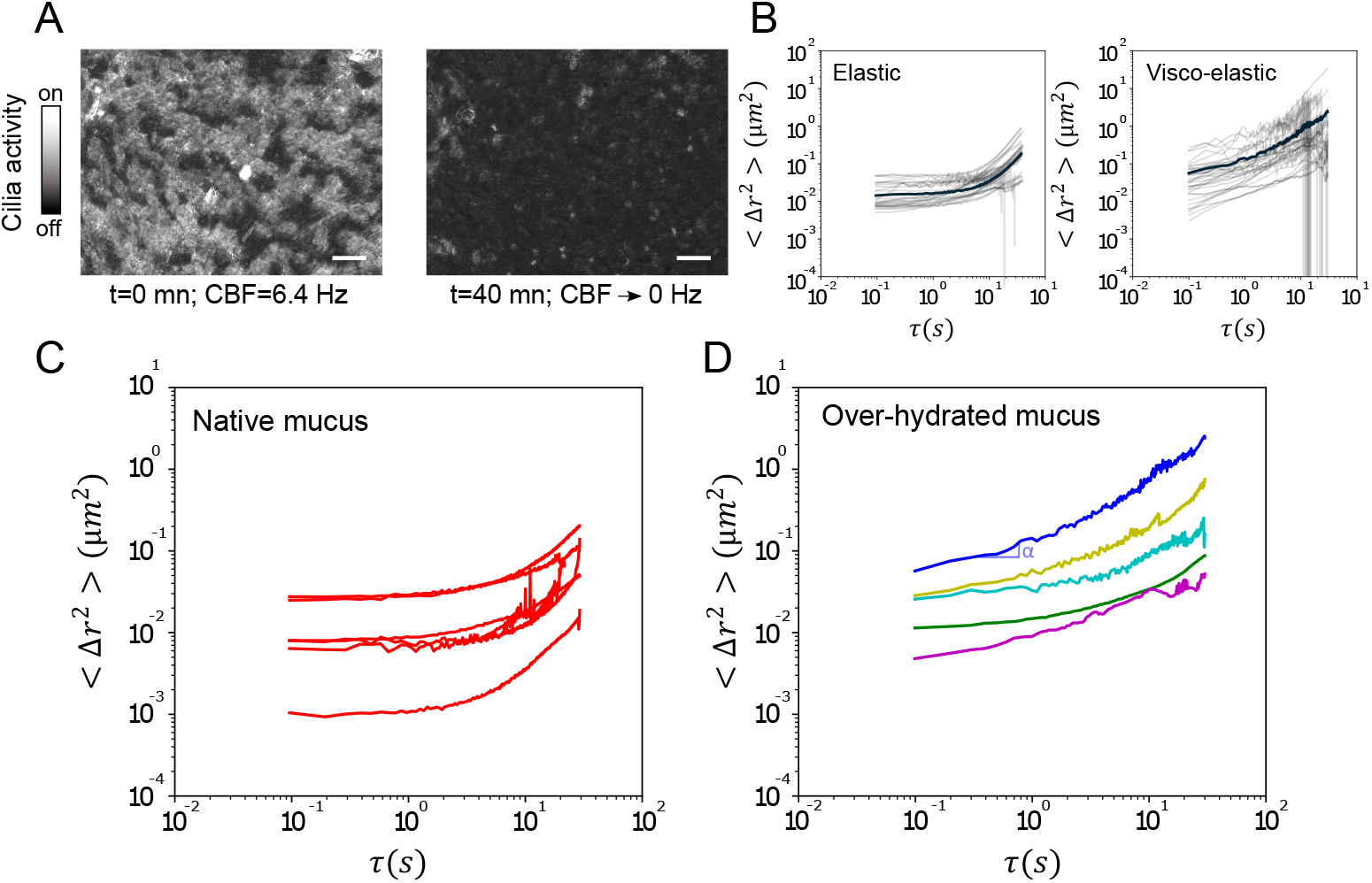
*In situ* microrheology via ciliary arrest. (A) Images showing the inhibition of ciliary activity over time after application of 22.5 mM cinnamaldehyde. Ciliary beat frequency (CBF) drops from 6.4 Hz (t=0) to 0 Hz (t=40 min). The images are the result of a standard deviation projection over 500 frames acquired at 25 fps. Scale bars: 20 µm. (B) Representative Mean-Square Displacement (MSD) curves from a single vortex (N=50 beads, light curves) and their average (dark curve) in the case of an elastic response (left) and a viscoelastic one (right). (C) MSD curves showing “native” elastic behavior, characterized by a plateau (*α* = 0). (D) MSD curves showing “over-hydrated” viscoelastic behavior, characterized by a slope (0 < *α* < 1). The exponent *α* is extracted from the MSD in the regime preceding the onset of drift (see Materials and Methods).

**Figure 3B** displays Mean Square Displacement (MSD) measurements obtained from the tracking of fluorescent beads in two distinct vortices. The dispersion of individual bead trajectories (light curves, N ≈ 50) indicates spatial heterogeneity. However, the consistent slopes across beads suggest a coherent local rheology, justifying the use of ensemble-averaged MSDs. Two distinct rheological behaviors emerged from these measurements. The first shows a clear plateau (slope *α* = 0) at short time (*τ* < 5 s), this is characteristic of an elastic solid. The height of this plateau is a direct measure of the constrained motion of the beads and is inversely related to the mucus stiffness (*G*′). At longer time (*τ* > 5 s), we observe a non-Brownian drift artifact (*α* = 2), likely due to sample drift. The second one, shows a consistent slope (0 < *α* < 1) across the entire time range, characteristic of a viscoelastic fluid.

When analyzing all 11 vortices (**Figures 3C and 3D**), we observed a correlation between the rheological behaviors and the hydration state of the mucus. The elastic, solid-like responses (**Figure 3C**) correspond exclusively to native mucus vortices, whereas the viscoelastic responses (**Figure 3D**) correspond to the “over-hydrated” conditions (regular addition of PBS). In the following, we leverage the diverse rheological behaviours to explore the link between mucus rheology and transport properties using the metrics (Q) previously defined.

### Efficiency of mucus transport does not correlate with its rheological properties

Building on the quantification of both mucus flow rate and rheological properties, for each mucus vortex, we investigated the relationship between mucus rheology and transport. We extracted, from microrheology experiments, the slope *α* of the MSD curves (**Supplementary Figure S1**) and plotted it as a function of flow rate (Q) (**Figure 4**). We found two distinct regimes. Vortices made of over-hydrated viscoelastic (*α* > 0) mucus exhibit flow rates above a critical threshold (*Q*_*c*_ ≈ 3 · 10^5^ *µ*m^3^/s, dark area). In contrast, vortices made of native mucus, elastic behaviour (*α* = 0), all exhibit a flow rate that falls below the critical threshold (*Q* < *Q*_*c*_, light area).

**Fig. 4.**
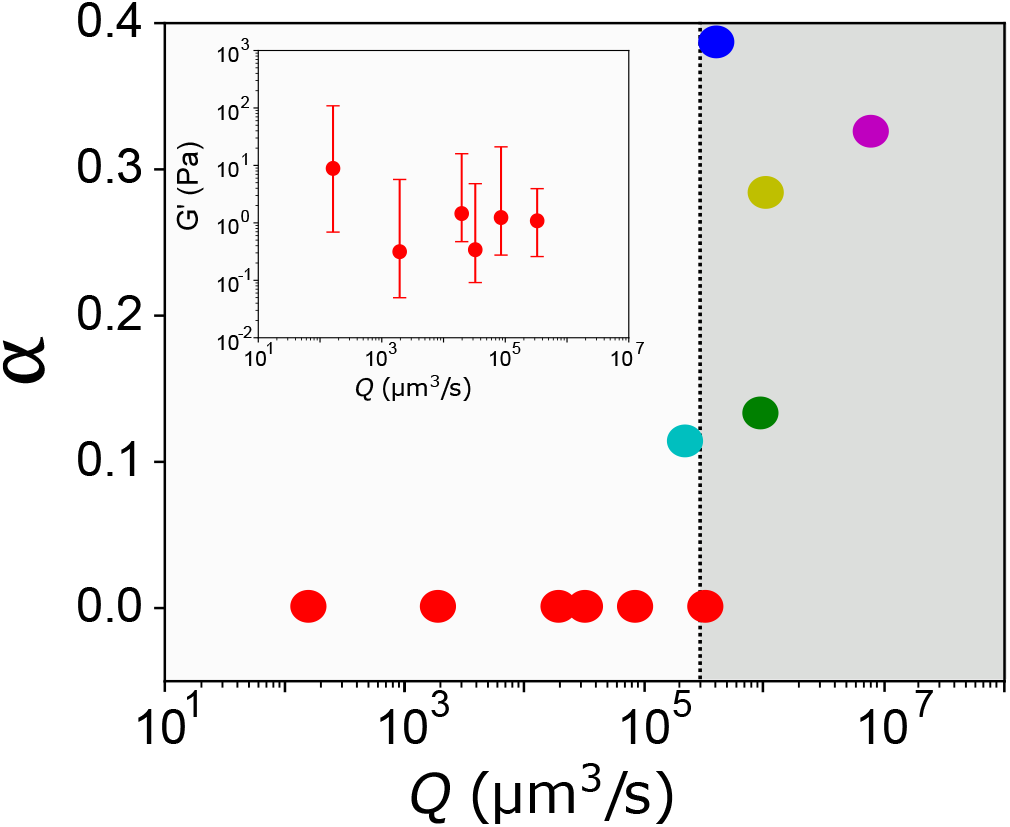
Connecting mucus rheology to transport efficiency. Exponent *α* plotted against flow rate Q for the 11 studied vortices. The data reveals two distinct transport regimes: an elastic regime (*α* = 0) at low Q (light area) and a viscoelastic regime (*α* > 0) at high Q (dark area). The transition occurs at *Q*_*c*_ ≈ 3 *·* 10^5^*µm*^3^/*s*. Inset : storage modulus *G*′ plotted against flow rate Q for samples within the elastic regime. Error bars represent the minimum and maximum *G*′ values obtained from microrheology, reflecting the spatial heterogeneity measured across all tracked tracer beads within each mucus sample.

In the high flow rate regime (*Q* > *Q*_*c*_, viscoelastic mucus) transport efficiency appears to saturate. Higher *α* (further fluidization) yields no proportional increase in flow rate. This indicates a decoupling between the rheological exponent *α* and transport efficiency (*Q*) in this regime.

For elastic mucus (*Q* < *Q*_*c*_), flow rates still span several orders of magnitude (10^2^ to 10^5^ µm^3^/s). We investigated the role of mucus stiffness in this variability. The storage modulus *G*′ was estimated from the MSD plateau using the generalized Stokes–Einstein relation. Plotting *G*′ against Q for these native mucus samples (**Figure 4, Inset**) shows that the ranges of *G*′ values (min–max across tracked beads) strongly overlap across samples displaying very different transport rates. Across the different experimental conditions, we computed the Spearman rank correlation coefficient between Q and the mean value of *G*′ for each sample. The analysis yielded *ρ* = − 0.37 (p = 0.47), indicating no detectable monotonic relationship between mucus stiffness and transport rate. Bulk elastic properties alone therefore cannot account for the observed transport behavior. This conclusion is further supported by experiments in which native mucus is replaced by PDMS: despite its markedly different bulk mechanical properties, transport remains efficient, indicating that transport is not controlled by bulk rheology but by the properties of the cilia–mucus interface.

In summary, while hydration determines the global transport and rheological regime (elastic for *Q* < *Q*_*c*_ vs viscoelastic for *Q* > *Q*_*c*_), the bulk stiffness (*G*′) of the native mucus does not regulate its transport efficiency. This result supports the existence of an interfacial layer that mechanically isolates the cilia from the bulk mucus.

In the next section, we use cilia as a physical probe to investigate the properties of the interface.

### Probing the cilia-mucus interface during transport slowdown

It is currently generally accepted that during the effective stroke, the ciliary tip penetrates the mucus by approximately 0.5 µm [47]. Cilia are therefore excellent candidates for probing the nature of the medium in which they beat. Analyzing the ciliary beat pattern can provide information on the local physical properties of the cilia-mucus interface. We developed optical flow algorithms to quantify the beat patterns of bundles of cilia (**Figures 5A & B**). **Figure 5C** shows the computed flow vectors over a representative analysis window. Each window was chosen to match the scale of a single cell’s apical surface—typically 50 µm^2^. These cellular-sized areas contain approximately 15 ciliary bundles beating in a coordinated fashion. For each selected area we extract the temporal evolution of the mean magnitude and the mean angle of the optical flow vectors. The trace of the temporal evolution of the velocity magnitude reveals the characteristic asymmetric beat pattern: sharp peaks correspond to the rapid forward stroke, while lower peak values correspond to the recovery stroke (**Figure 5D**). The velocity of the tip of a ciliary bundle during forward stroke (*v*_forward_) is approximated as the high peak magnitude. Simultaneously, the unwrapped angle, which represents the cumulative change in orientation, clearly shows stable plateaus for both stroke and recovery phases. It allows for the measurement of the forward stroke duration (*T*_forward_), representing 35-40% of the total cycle (see **Supplementary Figure S2**). The amplitude of a ciliary beat (**Figure 5I**) is estimated as the product *A* ≈ *v*_forward_ × *T*_forward_, and the Ciliary Beat Frequency (CBF) is derived from the power spectrum of the magnitude signal (**Figure 5E**).

**Fig. 5.**
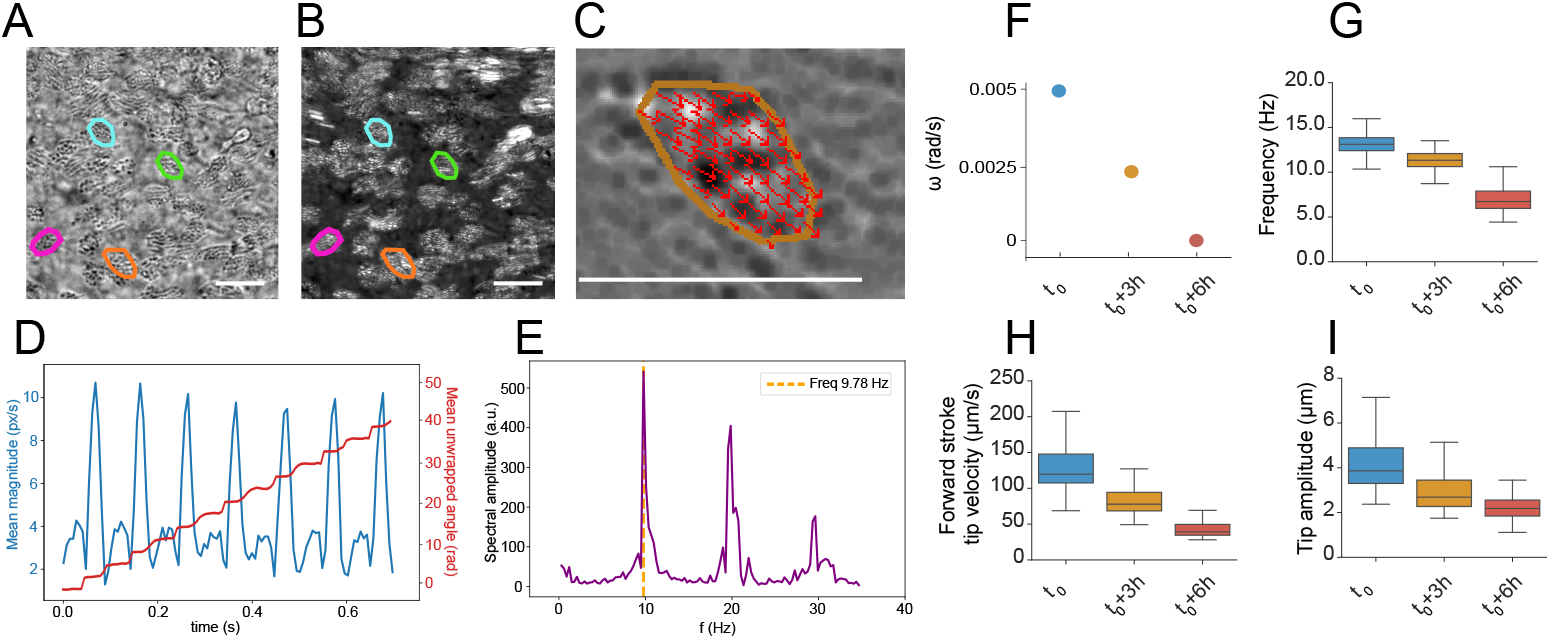
Evolution of ciliary beat pattern during mucus transport slowdown. (A, B) Examples of manually selected cellular-sized areas used for optical flow analysis. (C) Magnified view of a single cellular-sized area, overlaid with the computed optical flow vectors. Scale bars: 20µm. (D) Representative time traces extracted from the selected area (mean magnitude and unwrapped angle). (E) Power spectrum analysis used to determine the Ciliary Beat Frequency (CBF). (F-I) Kinematic parameters for a single vortex shown at three time points: *t*_0_ (rapid transport), *t*_0_ + 3*h* (slower transport), and *t*_0_ + 6*h* (stopped mucus). Data is compiled from ≈ 60 cellular-sized areas (each containing several cilia bundles) at each time point. (F) Vortex angular velocity (*ω*). (G) Ciliary Beat Frequency (CBF). (H) Forward stroke tip velocity (*v*_forward_). (I) Estimated tip amplitude.

To correlate these kinematic parameters – frequency, tip velocity, tip amplitude – with transport efficiency, we imaged both mucus transport and underlying cilia dynamics on a mucus vortex undergoing a decrease in angular velocity until arrest (**Figure 5F**). **Supplementary Video S5** illustrates progressive mucus slowdown and corresponding changes in ciliary beat patterns at 0 h, 3 h, and 6 h. Data compiled from approximately 60 cellular-sized areas over a time course of 6 hours, reveals a simultaneous decrease in all parameters, with relative variations of Δ*f*/*f* = 49%, Δ*v*/*v* = 67%, and Δ*A*/*A* = 46% (**Figures 5G–I**).

Under the assumption that cilia generate a constant forward force [48], this force must be balanced by the viscous drag at the interface. Consequently, the observed drop in velocity indicates a corresponding rise in the effective viscosity of the interfacial layer (*η*_interface_). This increasing viscosity is the mechanical signature of local dehydration. It suggests that transport failure occurs when the interfacial viscosity becomes too high for the cilia, slowing the beat until flow stops. To get better insight into the properties of the cilia-mucus interfacial layer responsible for mucus arrest, we used measured parameters of the beat patterns to inform a hydrodynamic model.

### Inferring the rheological properties of the cilia-mucus interface that leads to mucus arrest

We model the beating of a cilium immersed in a three-dimensional two-phase flow as previously described [29]. **Figure 6A-C** shows a schematic of the computational model. The cilium is modelled by an elastic filament actuated at its base. The basal angle *θ*(t) of the cilium varies sinusoidally to achieve a periodic beating. The two-phase flow, that describes the periciliary layer (PCL) and the surface fluid on top of cilia, is simulated using the Shan-Chen model in a lattice Boltzmann (LB) solver. In this framework, we vary two control parameters: the ciliary beat frequency and the viscosity ratio between the surface fluid and the PCL. Finally, the hydrodynamic coupling between the cilium and the surface fluid is as follows: the ciliary beating drives the fluid flow, and the fluid flow in turn affects the deformation of the cilium.

**Fig. 6.**
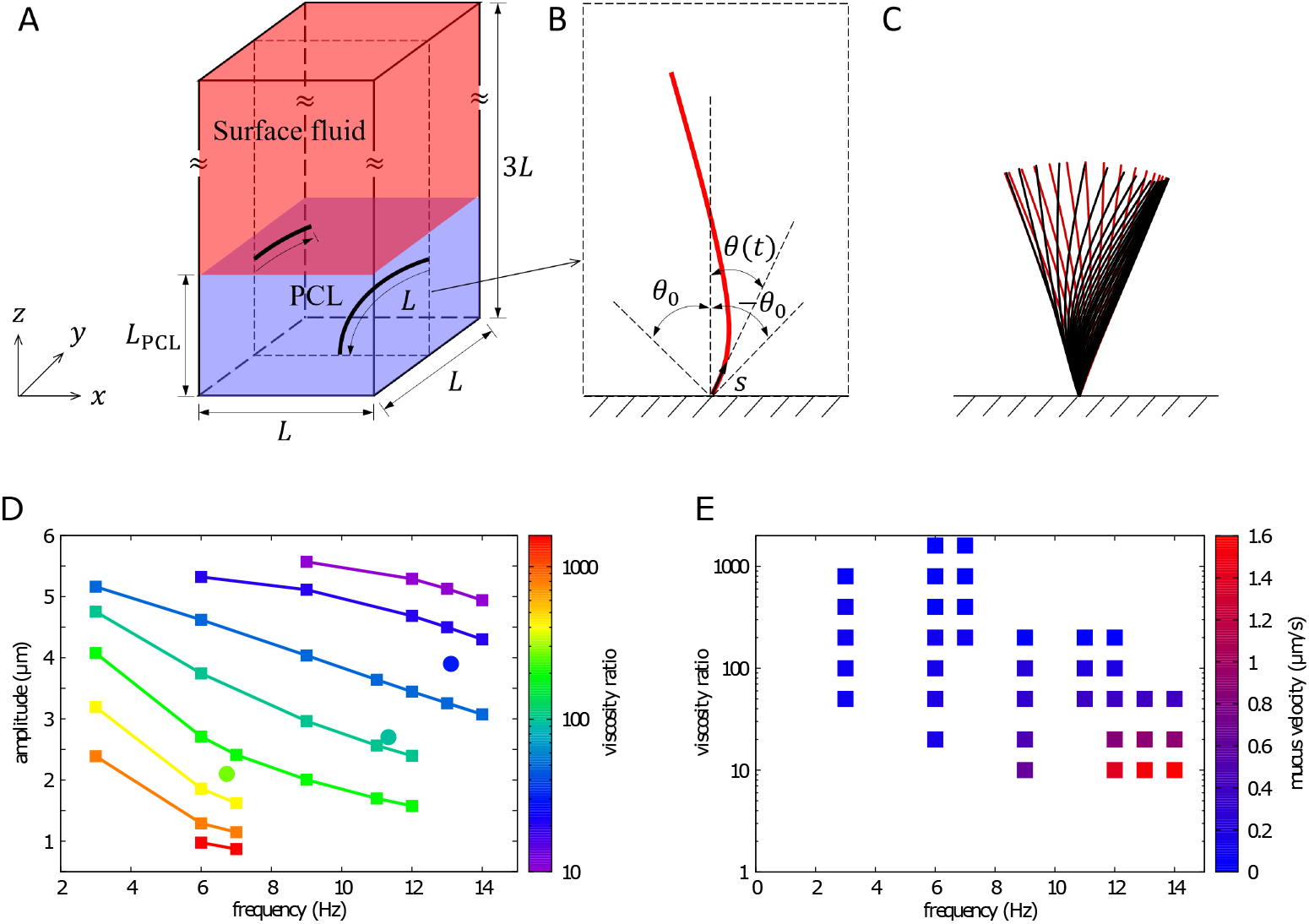
Hydrodynamic model of ciliary beating. (A) Schematic of the computational domain. (B) Schematic of a beating cilium actuated by its basal angle *θ*(*t*). (C) Beating pattern of a cilium. The positions of the cilium during the fast forward power stroke are represented in red, while the positions during the slow backward recovery stroke are in black. (D) Phase diagram of the viscosity ratio between the surface fluid and the PCL against beat amplitude and beat frequency. Square symbols are the simulated points and circle symbols are the experimental ones. The values of the viscosity ratio for experimental data points are inferred from the simulation. (E) Computed phase diagram of the velocity of the surface fluid against the viscosity ratio and the beat frequency.

Using this model, we investigated the effects of the variations of the viscosity ratio and the beat frequency on the amplitude of the beat pattern and on the fluid transport velocity. The resulting phase diagrams are shown on **Figure 6D & E**. When beat frequency and beat amplitude decrease simultaneously, the model predicts an increase in the viscosity ratio between the surface fluid and the PCL. This accurately recapitulates our experimental observations. Specifically, as the angular velocity of a mucus vortex declines over time due to dehydration of the cilia-mucus interface, we measured a decrease in both ciliary beat frequency and amplitude (**Figure 5F, G & I**).

By mapping the frequency and amplitude values corresponding to an arrested mucus onto the computed phase diagram, we can deduce the order of magnitude for the critical viscosity of the cilia-mucus interface at which mucociliary transport fails. For a measured frequency of 6.73 Hz and a beat amplitude of 2.1 µm, and assuming the PCL viscosity is comparable to that of water, we infer a critical interfacial viscosity of *η*_critical_ =296 mPa · s.

The model also enables direct prediction of the surface fluid velocity, associated to a viscosity ratio and a cilia beat frequency. The corresponding phase diagram is shown in **Figure 6E**. To compare a numerically predicted velocity with an experimental measurement, we need to normalize the in-silico cilia density to match the experimental cilia density. The simulation domain contains a single ciliated element. The horizontal dimensions of the simulation box are equal to the length of a cilium, L=7 µm. Experimentally, a surface area of 7×7 µm^2^ corresponds to the apical surface of a single ciliated cell, which contains an average of 15 bundles of cilia. Consequently, the computed velocities must be multiplied by a factor of 15 to allow for a direct comparison with experimental values (in a low Reynolds regime, velocities are additive). Following the normalization of ciliary density, for an experimentally measured beat frequency f=13 Hz and a beat amplitude A=3.9 µm, the model predicts a viscosity ratio of 34 and a surface fluid velocity of 10 µm/s. This predicted velocity is consistent with the characteristic mucus velocities observed under hydrated conditions. Conversely, for the critical case where f=6.7 Hz and A=2.1 µm, corresponding to a viscosity ratio of 296, the predicted flow velocity at the cilia-mucus interface drops to 0.5 µm/s. This result is qualitatively consistent with the observation of an arrested mucus.

## 3 Discussion

The traditional paradigm in the context of chronic respiratory research assumes a direct causal link between the bulk rheological properties of mucus and transport efficiency, where increased viscosity and elasticity are the primary drivers of clearance failure. However, our results demonstrate a fundamental decoupling between bulk rheology and transport. The most striking evidence is the observation that cilia can efficiently propel cross-linked PDMS, an elastomer with an elastic modulus (*G*′ ∼ 10^6^Pa) six orders of magnitude stiffer than native mucus (*G*′ ∼ 1Pa) at physiological velocities. These findings, corroborated by the lack of correlation between the storage modulus (G’) and volumetric flow rate (Q) in native mucus, indicate that bulk rheology is not the primary limiting determinant of ciliary propulsion.

Instead, we propose that mucociliary clearance is governed by the hydration state of a thin layer at the cilia-mucus interface. As the interface dehydrates, the effective viscosity (*η*_interface_) increases. Based on our hydrodynamic model and the assumption of a constant ciliary forward force, the increase of viscosity results in an increase of the viscous drag until it overcomes the ciliary motor capacity. This leads to the observed simultaneous decrease in ciliary beat frequency, tip velocity, and amplitude during transport slowdown. Mucus arrest occurs for a critical value of *η*_interface_ ∼ 300 mPa · s. This value is consistent with measurements previously done by Hill et al. [48]. Using magnetic tweezer, they measured the force exerted by cilia and deduced a critical viscosity of 100 mPa · s.

A striking observation is the high sensitivity of the system to its hydration state. Specifically, the fact that the angular velocity of mucus vortices continuously decreases even when the sample is maintained in a humidity-saturated environment suggests that very small variations in the regulation of the cilia-mucus interface hydration are responsible for the significant disparities in transport efficiency observed in these systems. In air liquid interface cultures, we cannot exclude the possibility that certain ion channels such as CFTR or ENaC, which are critical regulators of periciliary layer (PCL) hydration, may not be fully functional [32]. More broadly, the criticality of hydration must be considered within the context of climate change, where chronic exposure to arid environments represents an escalating respiratory risk [49]. This challenge similarly arises during mechanical ventilation in intensive care; while it is essential that ventilatory gases are warmed and humidified, a precise balance is required, as excessive humidity can overwhelm mucociliary transport and increase the mucosal burden [50].

These findings may have significant clinical implications. They suggest that arrest of mucus might result from interfacial failure due to local dehydration rather than the composition of mucus itself. Consequently, therapeutic strategies might be more effective if they prioritize the maintenance of the hydration of the cilia-mucus interface rather than simply targeting the breakdown of mucus mesh with mucolytic agents. Finally, this ALI model provides a quantitative, standardized assay for testing the effect of drugs on the mucociliary clearance efficiency, provided that we account for the critical role of interfacial hydration to ensure reproducible measurements and quantitative comparison between samples.

## 4 Methods

### Cell Culture

Reconstituted human bronchial epithelium cultures (MucilAir™) were obtained from Epithelix Sàrl (Switzerland) and cultured at the Air-Liquid Interface (ALI). Cultures were maintained at 37°C in a humidified incubator with 5% CO_2_, with the culture medium (MucilAir™ culture medium) replaced every two days. Two types of cultures were used depending on the experiment. For mucus transport and rheology quantification, we used fully differentiated cultures. Mucus vortices typically formed 2–3 days following an initial apical wash. To maintain stable vortex rotation, a minimal volume of PBS (1.5 *µ*L) was added apically every day, ensuring surface hydration without flooding the interface. For the analysis of ciliary beat patterns, we used partially differentiated cultures. These samples exhibited a lower density of cilia (≈ 40% coverage vs. ≈ 70% in fully differentiated inserts), which facilitated the optical resolution of the beating kinematics. The same daily hydration protocol was applied to induce and stabilize mucus vortices in these samples.

### Transport Quantification

Mucus transport was imaged using bright-field microscopy on an inverted Nikon Eclipse Ti microscope equipped with a × 20 objective and a Lumenera Infinity camera. Environmental conditions were maintained at 37°C with a humidified atmosphere containing 5% CO_2_. To quantify transport, cellular debris naturally trapped in the mucus were manually tracked to determine their trajectories. From these, we computed the angular velocity *ω* (averaged over at least 5 trajectories per vortex) and estimated the vortex radius *R*. The mucus height *h* was measured using the calibrated Z-stage of the microscope, defined as the vertical distance between the apical cell surface and the highest focused debris. Finally, the volumetric flow rate was calculated as 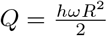.

### Cilia Inhibition

Ciliary beating was temporarily inhibited using cinnamaldehyde (Sigma-Aldrich). The inhibitor was added to the basal medium at a final concentration of 22.5 mM, 40 minutes prior to observation. The total cessation of ciliary activity was systematically confirmed via video microscopy before any further measurements.

### Passive Microrheology

Passive microrheology was performed using 200 nm diameter fluorescent carboxylate-modified polystyrene beads (Invitrogen). To prevent non-specific interactions with the mucus mesh, beads were coated with polyethylene glycol (PEG) following the protocol described by Schuster et al. [15]. The tracers were diluted in PBS and added to the apical surface during the hydration step. Particle tracking was performed on a Leica MICA microscope in widefield mode using a × 63 water immersion objective. Acquisitions were recorded for 2 minutes at 10 fps to ensure statistical robustness for low-frequency analysis. The Brownian motion of 30–50 beads per vortex was analyzed using a custom Python script based on the *trackpy* library. The Mean-Square Displacement (MSD, ⟨Δ*r*^2^⟩) was calculated from particle trajectories. With a spatial resolution of 0.09 µm/pixel and a tracking precision of *σ*_track_ = 0.1 pixel, our theoretical noise floor is 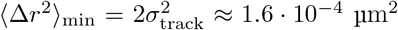 [51]. All measured MSD data were significantly above this threshold, confirming that the observed displacements represent physical particle motion rather than tracking artifacts. The MSD was analyzed in log–log representation. The onset of drift was detected by a relative increase of more than 35% in the local logarithmic slope. The exponent *α* was obtained from a linear fit of log(MSD) versus log(*τ*) restricted to the regime preceding this deviation. Trajectories with *α* < 0.1 were considered to display an elastic plateau (*α* ≈ 0). In this case, the storage modulus *G*′ was estimated using the generalized Stokes–Einstein relation [43], 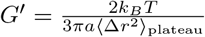, where *a* is the bead radius.

### Ciliary Beat Pattern Analysis

Ciliary beating was recorded at 148 fps using high-speed video microscopy on an inverted Nikon Eclipse Ti microscope equipped with an Andor Sona sCMOS camera. We used a water immersion objective (× 60 WI, NA 1.20) with a long working distance (0.28–0.31 mm) to image through both the transwell membrane and the epithelial layer. During acquisition, the transwell insert was placed directly on a glass coverslip. A droplet of culture medium was added between the coverslip and the membrane to ensure liquid contact and nutrient supply. The system was enclosed in a custom 3D-printed chamber to maintain a sterile environment. Stable conditions (37°C, 5% CO_2_) were maintained using a humidified airflow delivered to the stage. Quantitative analysis of ciliary motion was performed using a custom Python script based on dense optical flow. Local velocity vector fields were computed for each pixel using the Farnebäck algorithm (OpenCV library). A window size of 15 pixels, approximately corresponding to the size of ciliary bundles, was selected. Functionally, this window defines the local image region over which intensity patterns are correlated between consecutive frames to estimate a displacement vector. A three-level pyramidal decomposition, in which optical flow is first computed on downsampled images and progressively refined at full resolution, was applied to ensure robust estimation of large displacements between consecutive frames. Optical flow was specifically analyzed within manually defined regions of interest (ROIs), typically corresponding to the size of individual ciliated cells. Representative optical flow vector fields overlaid on ciliary motion are shown in **Supplementary Video S5**. From the resulting flow fields, velocity magnitude and orientation signals were extracted and averaged over each ROI. The ciliary beat frequency (CBF) was obtained from the velocity magnitude using power spectral density analysis. Beat asymmetry was quantified from the continuous unwrapped angle of the flow vectors, which converts the cyclic angular signal—characterized by artificial jumps between -180° and +180°—into a continuous time series by cumulatively integrating successive angular variations over time. This enabled identification of directional turnover points within each beat cycle and precise measurement of the forward (*T*_forward_) and recovery (*T*_recovery_) stroke durations, as well as beat amplitude (**Supplementary Figure S2**).

### Numerical simulations

The beating of a cilium in a three-dimensional two-phase flow is modeled based on a previously established approach [29]. A schematic of the computational model is shown in **Figure 6A**. The cilium is represented as an elastic filament actuated by a sinusoidally varying basal angle *θ*(*t*) as shown in **Figure 6B**, and its dynamics are governed by the equations of elasticity with a time-dependent bending rigidity *B*. The filament is stiff during the forward power stroke (*B*_max_ = 1 × 10^−23^ N m^2^ [52]). Upon entering the backward recovery stroke, it softens (*B* decreases to *B*_max_/*r*_*B*_ with *r*_*B*_ = 70), after which its stiffness gradually recovers to *B*_max_. This strategy reproduces the essential feature of asymmetric ciliary beating [29]. The maximum amplitude of the basal angle is set to *θ*_0_ = *π*/9. The ratio between the durations of the forward power stroke and the backward recovery stroke is fixed at 1/2, consistent with characteristic values measured *in vitro* [53, 54]. The resulting motion yields a two-dimensional beating pattern that consists of a fast power stroke with an elongated shape and a slow recovery stroke with a bent shape as illustrated in **Figure 6C** and **Supplementary Video S6**. The surrounding two-phase flow is resolved using the Shan–Chen model within a lattice Boltzmann framework, which enables stable phase separation and allows the fluid–fluid interface to evolve dynamically. In **Figure 6A**, the periciliary layer (PCL) and the surface fluid are indicated in blue and red, respectively. The thickness of the PCL is set to *L*_PCL_ = 0.8*L* to allow the cilium to slightly penetrate into the surface fluid, where *L* denotes the length of the cilium. A no-slip boundary condition is applied at the bottom boundary of the computational domain, while a free-slip boundary condition is imposed at the top boundary. Periodic boundary conditions are prescribed on the remaining four boundaries. The computational domain is chosen sufficiently large to minimize boundary effects on the ciliary beating dynamics. Finally, the two-way coupling between the cilium and the surrounding fluid is handled using the immersed boundary method, whereby the ciliary beating drives the fluid flow and the fluid motion in turn affects the deformation of the cilium.

## Supporting information

Supplementary Informations

## Acknowledgements

This work was supported by the BonchoClogDrain project (ANR-22-CE30-0045) funded by the French National Research Agency (ANR)

