## Supplementary Informations for "Disentangling mucus rheology and transport efficiency in human airways"

### Supplementary Information for:
Disentangling mucus rheology and transport efficiency in human airways

Alice Briole, Qian Mao, Umberto D'Ortona, Julien Favier,
Annie Viallat, Etienne Loiseau

#### Supplementary videos

**Video S1: Transport of a PDMS object.** Bright-field microscopy showing the transport of a cross-linked PDMS elastomer placed on a mucus-free epithelial surface. Following a slight apical rehydration (1.5 µL), the PDMS object is set into rotational motion by ciliary beating.

**Video S2: Macroscopic mucus transport in a rotating vortex.** Bright-field microscopy showing the rotation of a mucus vortex. Cellular debris maintain fixed relative positions during rotation, demonstrating that the mucus layer is transported as a cohesive solid body.

**Video S3: Comparison of native and over-hydrated mucus transport.** Side-by-side comparison of mucus transport and aspect. *Left panel:* Native condition. The mucus is concentrated. *Right panel:* Over-hydrated condition (following regular PBS additions). The mucus appears smoother.

**Video S4: Inhibition of ciliary beating over time.** Three-panel view showing the progressive arrest of ciliary activity at $t = 0$, $t = 20$, and $t = 40$ minutes after basal addition of 22.5 mM cinnamaldehyde. This sequence demonstrates the complete cessation of beating required for passive microrheology measurements.

**Video S5: Optical flow analysis of ciliary beat patterns.** The *left column* shows mucus vortices at 0 h, 3 h, and 6 h, illustrating the progressive slowdown of transport. The *right column* displays representative ciliary beat patterns from the corresponding time points, overlaid with optical flow vectors within selected regions of interest (ROIs). Vectors indicate the magnitude and direction of the local velocity field in real time.

**Video S6: Numerical simulation of ciliary beating.** Numerical simulation of a beating cilium with the beating frequency set to 9 Hz and a viscosity ratio of 10.

### Supplementary figures

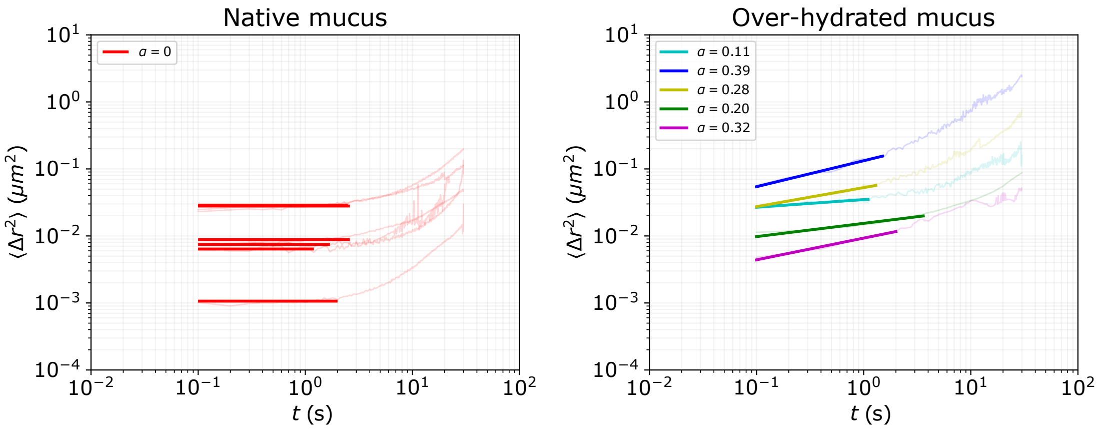


Figure S1: **Determination of the exponent $\alpha$ from MSD curves.** MSD curves shown in the main text with the corresponding linear fits in log–log scale and the corresponding $\alpha$ value. The fitting interval is restricted to the initial regime preceding the drift onset, as defined in Materials & Methods.

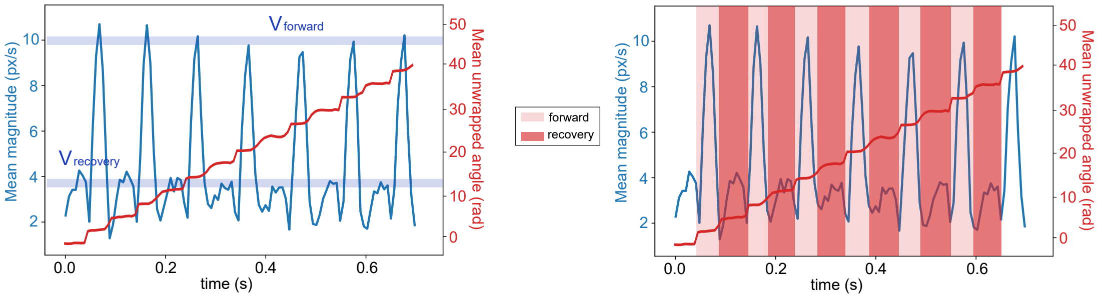


Figure S2: **Method for extracting kinematic parameters from optical flow.** Both panels display the superimposed time evolution of the velocity magnitude and the unwrapped angle measured on a cellular-sized area. *Left panel.* Quantification of stroke velocities. Horizontal lines indicate the extracted forward stroke velocity (corresponding to high-magnitude peaks) and the recovery stroke velocity (corresponding to low-magnitude peaks). *Right panel.* Quantification of forward and recovery stroke durations. Colored time intervals, whose boundaries are defined by inflection points of the angular signal, are used to extract the forward ($T_{\text{forward}}$) and recovery ($T_{\text{recovery}}$) durations. The beat amplitude is estimated as the product of the forward stroke velocity and duration ($A \approx V_{\text{forward}} \times T_{\text{forward}}$)
